# Explainable Sleep Stage Classification with Multimodal Electrophysiology Time-series

**DOI:** 10.1101/2021.05.04.442658

**Authors:** Charles A. Ellis, Rongen Zhang, Darwin A. Carbajal, Robyn L. Miller, Vince D. Calhoun, May D. Wang

## Abstract

Many automated sleep staging studies have used deep learning approaches, and a growing number have used multimodal data to improve their classification performance. However, few studies using multimodal data have provided model explainability. Some have used traditional ablation approaches that “zero out” a modality. However, the samples that result from this ablation are unlikely to be found in real electroencephalography (EEG) data, which could adversely affect the importance estimates that result. Here, we train a convolutional neural network for sleep stage classification with EEG, electrooculograms (EOG), and electromyograms (EMG) and propose an ablation approach that replaces each modality with values that approximate the line-related noise commonly found in electrophysiology data. The relative importance that we identify for each modality is consistent with sleep staging guidelines, with EEG being important for most sleep stages and EOG being important for Rapid Eye Movement (REM) and nonREM stages. EMG showed low relative importance across classes. A comparison of our approach with a “zero out” ablation approach indicates that while the importance results are consistent for the most part, our method accentuates the importance of modalities to the model for the classification of some stages like REM (p < 0.05). These results suggest that a careful, domain-specific selection of an ablation approach may provide a clearer indicator of modality importance. Further, this study provides guidance for future research on using explainability methods with multimodal electrophysiology data.

## I. Introduction

Many methods have been developed for automated sleep staging in recent years. Most have used electroencephalograms (EEG) [1] or electrooculograms (EOG) [2], and only a few have utilized multimodal data [3][4]. Clinicians typically use multimodal data when scoring sleep stages, so the use of multimodal data, like EEG, EoG and electromyograms (EMG), might provide additional insight that could not be gained from unimodal data alone, as multimodal data facilitates more intricate recognition of human activity [4].

Moreover, while many studies have used deep learning (DL) for automated sleep staging, most of them do not give insight into the inner mechanisms of their classifiers [5]. DL models’ black-box nature may be problematic for clinical implementation because they are difficult for clinicians to interpret. of the papers that offer explainability, most have been developed for EEG [6]-[8]. Studies on multimodal data have, with a few exceptions [5],[6], not used explainability methods or provided insight into the importance of the modalities that they analyzed.

Several studies have used explainable artificial intelligence methods for classification with multimodal datasets [10] [11]. These studies use approaches like layer-wise relevance propagation (LRP) and ablation [10]. Ablation involves the removal of each modality and the calculation of the effect that its removal has upon the performance of the classifier. Like similar methods that perturb portions of the data, ablation methods can create new samples that are outside the data distribution upon which the classifier was trained and potentially give an inadequate explanation of the model [12]. While ablation approaches are simple and intuitive, it is important to consider the potential problems that could arise when they are applied in domains, like electrophysiology classification, for which they were not originally designed.

Existing multimodality studies that have used ablation have predominantly replaced each modality with zeroes [10][11]. While this “zeroing out” of a modality may not force a sample outside the data distribution if an appropriate data normalization method (i.e., z-scoring) is used, the resulting samples will nevertheless be unrealistic and will not align with how real-life samples appear.

To remedy this problem, here we present a method for automated sleep staging with multimodal electrophysiology data by adapting a 1-dimensional (1D) convolutional neural network (CNN) architecture originally developed for EEG-based sleep stage classification [13]. We also provide a novel approach for gaining insight into the relative importance of each modality to the classification of each sleep stage. our ablation approach seeks to ablate each modality in a way that creates new samples that (1) a classifier might encounter during a clinical implementation and (2) that would not be as distinct from the original data as “zeroed out” samples. We do this by replacing each modality with a sinusoid and Gaussian noise that mimics the line-related noise that is commonly found in electrophysiology recordings. We further compare the results of our explainability method with that of the traditional “zeroing out” ablation approach.

## II. Methods

Here we describe our study approach. We train a 1D CNN for sleep stage classification with multimodal data and output ablation-based explanations for insight into the modalities critical for classifying each stage. We use statistical tests to compare our approach with an existing method.

### A. Description of Data

We used sleep telemetry data from the PhysioNet [14] Sleep-EDF Database [15]. The data has been used in sleep stage classification studies in the past [16]. The dataset consists of 44 full night (approximately 9 hour) recordings collected from 22 healthy subjects. Each subject had two recordings: one following the administration of a placebo and one following the administration of temazepam. The data consisted of three electrophysiology modalities: EEG, EoG, and EMG. All modalities were recorded at a sampling rate 100 Hertz (Hz). For EEG, we used the FPz-Cz electrode. A marker recorded at 1 Hz indicated whether the sleep telemetry system was operating correctly, and a polysomnogram consisting of Awake, Movement, rapid eye movement (REM), Non-REM 1 (NREM1), NREM2, NREM3, and NREM4 periods was also included.

### B. Description of Preprocessing

We divided the data into non-overlapping, 30-second segments and extracted labels from polysomnograms at 30-second intervals. NREM3 and NREM4 stages were combined into a single NREM3 stage [17] and Movement samples were discarded. We discarded all samples that coincided with a telemetry device error. We applied z-score normalization for each individual modality within each recording. After segmentation, the dataset had 42,218 samples across all 22 subjects. Approximately 9.97% (4,213 samples), 8.53% (3,603 samples), 46.8% (19,755 samples), 14.92% (6,298 samples), and 19.78% (8,349 samples) belonged to the Awake, NREM1, NREM2, NREM3, and REM classes, respectively.

### C. Convolutional Neural Network

We adapted a CNN architecture initially developed for EEG sleep stage classification [13]. The architecture is shown in Figure 1 and was implemented with Tensorflow and Keras. When training the model, we used 10-fold cross-validation in which 17, 2, and 3 subjects in each fold were randomly assigned to training, validation, and test groups, respectively. While training the classifier, we used categorical cross entropy loss and weighted the loss for each class to account for class imbalances. We used the Adam optimizer [18] with an adaptive learning rate that decreased after every 5 epochs with no increase in validation accuracy. The optimizer used an initial learning rate of 0.001. During testing of each fold, we used the model weights from the epoch with the best validation accuracy. To account for class imbalances when assessing test performance, we calculated the precision, recall, and F1 score for each class in each fold. After computing the precision, recall, and F1 scores, we calculated their mean and standard deviation (SD) across all folds.

**Figure 1.**
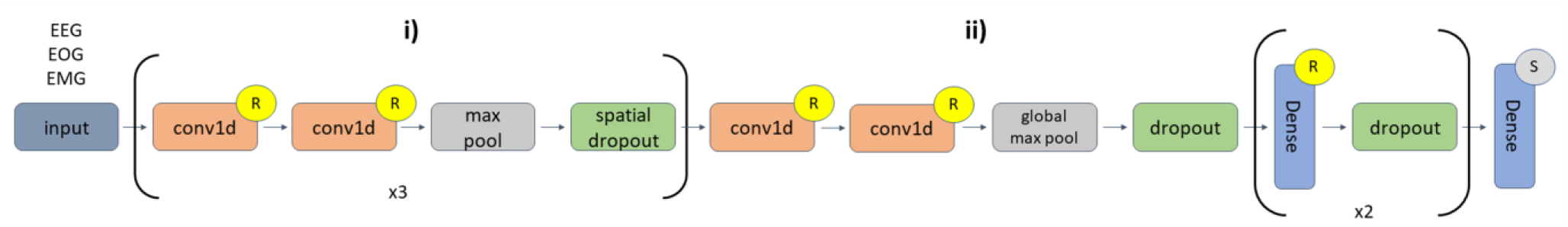
CNN Architecture. In the repeated layer **i)** of the diagram there are 6 1D convolutional (conv1d) layers. The first two conv1d layers have 16 filters with a kernel size of 5 followed by a max pooling layer with a pool size of 2 and a spatial dropout layer with a rate of 0.01. The second two conv1d layers (in **i**) had 32 filters with a kernel size of 3, followed by a max pooling layer with a pool size of 2 and a spatial dropout layer with a rate of 0.01. The third pair of conv1d layers (in **i**) have 32 filters with a kernel size of 3 followed by max pooling with a pool size of 2 and spatial dropout with a rate of 0.01. The last two conv1d layers (in **ii**) have 256 filters with a filter size of 3 followed by global max pooling and dropout with a rate of 0.01. The first dense layer has 64 nodes with dropout layer with a rate of 0.1. The second dense layer has 64 nodes with a dropout layer with a rate of 0.05. The last dense layer has 5 nodes. Layers with an “R” or an “S” indicate that they are followed by a ReLU or Softmax activation function, respectively.

### D. Ablation-based Global Explainability

We applied an ablation approach for insight into the importance of each modality to the identification of each sleep stage. Instead of zeroing out each modality, we replaced them with values that might be expected in electrophysiology data. Line-related noise often appears in electrophysiology data at around 50 Hz or 60 Hz as a result of the presence of power lines, lights, and other electronics near recording devices, and when an electrode is not working properly, it is common to find only line-related noise in that particular channel. For a sampling rate of 100 Hz, aliased 60 Hz noise should appear at around 40 Hz. As such, for our study, we replaced each modality with a combination of a 40 Hz sinusoid with an amplitude of 0.1 and Gaussian noise with a mean of 0 and SD of 0.1. Before ablation, we measured the weighted F1 score across all classes and the F1 scores for each individual class. We then ablated each modality and calculated the resulting change in performance (original F1 - New F1). We performed the ablation for each fold individually. For a comparison, we used a “zeroing-out” ablation approach [10] and performed two-tailed t-tests to compare the change in F1 associated with each class and modality for the two methods.

## III. Results and Discussion

In this section, we detail and discuss model performance and the insights gained by comparing the ablation methods.

### A. Model Performance Results

Table 1 shows the model’s test performance across all 10 folds. The classification of NREM1 samples obtained the lowest level of performance across all metrics, which is unsurprising given that the NREM1 stage was the smallest class. However, while the Awake and NREM1 classes had a comparable number of samples, the models obtained much higher performance for the Awake class. Additionally, the classifier obtained higher precision for the Awake class than all other classes except for NREM2, higher recall than all other classes except for NREM3, and a higher F1 score than all other classes except for NREM2. This makes sense given that EEG, EoG, and EMG Awake activity is very different from NREM activity and that EMG Awake activity is very different from EMG REM activity [17]. Additionally, the model obtained highest precision and F1 scores for the NREM2 class, which is reasonable given that nearly half of the dataset was composed of the NREM2 class.

**TABLE I.** Classification Performance Results

|  | Awake | NREM1 | NREM2 | NREM3 | REM |
| --- | --- | --- | --- | --- | --- |
| **Precision** | 72.25±07.12 | 36.20±03.98 | 79.35±03.92 | 56.78±18.35 | 69.04±07.14 |
| **Recall** | 70.90±07.02 | 46.28±13.52 | 68.71±08.51 | 78.22±10.24 | 63.26±06.69 |
| **F1** | 71.25±05.15 | 39.86±07.19 | 73.28±04.76 | 64.15±15.25 | 65.92±06.28 |

### B. Ablation-based Explainability Results

Figure 2 shows the explainability results for each method, along with the results of the statistical analysis. Panel A shows the change in F1 across all classes, and Panels B through F show the change in F1 for each individual class. For the line noise-related ablation analysis with the F1 score calculated for all classes, EEG was the most important modality by far, with a median reduction in F1 of nearly 60% following the ablation of EEG. It should be noted, however, that this change in F1 for all classes is skewed by the NREM2 and NREM3 classes. For the NREM2, NREM 3, and Awake classes, EEG was by far the most important modality while EoG and EMG had relatively little effect. EoG and EMG had larger effects upon the F1 score for the NREM1 and REM classes. For the NREM1 class, EEG and EoG had comparable effects upon the F1 score, though EEG had a slightly higher effect. Interestingly, the ablation of EMG for the NREM1 class seemed to have a beneficial effect, increasing the F1 score by as much as 6-7%. For the REM class, both EEG and EoG had a significant effect upon the F1 score, though EEG had a markedly larger effect. Additionally, for the REM class, EMG had an effect upon the F1 score as high as 15-16%. Also, given that our architecture was originally designed for EEG classification, it is possible that it didn’t effectively extract EMG features. This could explain the relatively low importance of EMG to the model.

**Figure 2.**
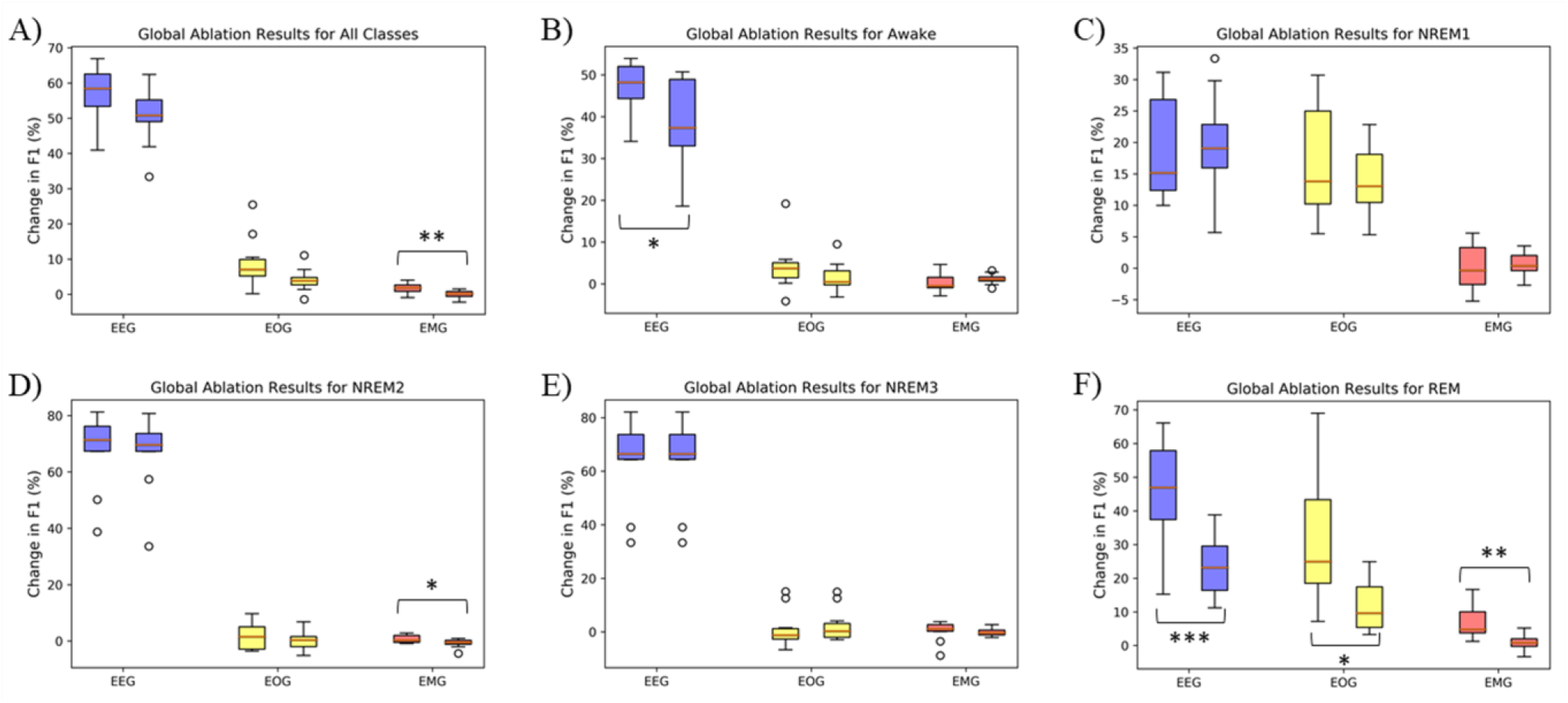
Results for Explainability Methods and Statistical Analysis. Panel A shows the explainability results across all classes, and Panels B through F show the results for individual classes. Blue, yellow, and red bars show the results for EEG, EOG, and EMG ablation, respectively. The leftmost of each pair of boxes shows the results for our ablation approach, and the rightmost shows the results for zeroing out each modality. The y-axes show the percent change in F1 following ablation, where positive and negative changes indicate decreases and increases in the F1 score, respectively. Some pairs of boxes are accompanied by *, **, or ***, which correspond to a two-tailed t-test p-value less than 0.05, 0.01, and 0.001, respectively.

The results of zeroing out each modality were highly similar to the results of applying line-related noise ablation. Most pairs of F1 ablation scores did not have a significant difference between them. However, when including all classes, EMG importance was significantly different between ablation methods (p < 0.01). For individual classes, the EEG of the Awake class (p < 0.05) and the EMG of the NREM2 class (p < 0.05) had significant results. Interestingly, all modalities had significant differences for the REM category (p < 0.05). In general, when a significant difference existed between importance assigned to a modality by the two explainability methods, our line-related noise ablation method seemed to give greater importance to the modality. This may indicate that our approach accentuates the importance of some modalities, which could be attributed to its use of ablation values that are more similar to real data.

The results of both methods generally fit with sleep scoring guidelines [17]. It was expected that EEG would be most important given that there is significant variation in EEG between NREM, Awake, and REM stages [17]. While relying upon EEG would enable the classifier to obtain good differentiation between Awake/REM and NREM, it is likely that relying upon EEG would not help with good classification between Awake and REM to the same degree. It would be logical for EMG to be important for classifying Awake and REM samples, as more movement might be expected in Awake than in NREM and as REM EMG activity would be much less than Awake and NREM activity [17]. It is interesting that EEG and EOG had comparable importance for NREM1 classification. Given that the classifier obtained lowest performance for the NREM1 class, the classifier may have inappropriately relied upon EOG for the identification of samples belonging to that class.

### D. Future Work

Examining model architectures that might better extract EMG features could be beneficial. While our explainability results fit with sleep scoring guidelines, suggesting the broader generalizability of the classifier, there are still areas in which our explainability could potentially be improved. For our ablation method, we sought to provide an alternative to traditional ablation approaches that zero out a feature by instead replacing the modalities with values similar to artifacts that naturally occur in electrophysiology recordings. When we approximated line-related noise, we had two parameters: the amplitude of the sinusoid and the SD of the Gaussian noise. It is possible that exploration of additional parameter values might improve explanations. Also, exploring explainability methods that do not require ablation or perturbation of features could further improve explanations.

### A. Conclusion

In this study, we trained a classifier on multimodal data for sleep stage classification. We further proposed a novel ablation-based approach that sought to provide more realistic ablated samples than methods that simply zero out a particular modality. A comparison of our method with the traditional ablation approach indicated that the importance values were comparable with the exception of a few instances in which our method seemed to accentuate the importance of some modalities. More broadly, this work has implications for ablation-based explainability in other data types, as it suggests that more careful consideration of how features are ablated in light of their domain may provide increases in importance metrics and clarify the relative importance of features.

## Acknowledgment

We thank Felipe Giuste and Wenqi Shi for their advice and assistance with computational resources. We thank Mohammad Sendi for helping edit the paper.

## References

[1] A. Sors, S. Bonnet, S. Mirek, L. Vercueil, and J. F. Payen, “A convolutional neural network for sleep stage scoring from raw single-channel EEG,” Biomed. Signal Process. Control, vol. 42, pp. 107–114, 2018, DOI: 10.1016/j.bspc.2017.12.001.

[2] M. M. Rahman, M. I. H. Bhuiyan, and A. R. Hassan, “Sleep stage classification using single-channel EOG,” Comput. Biol. Med., vol. 102, no. June, pp. 211–220, 2018, DOI: 10.1016/j.compbiomed.2018.08.022.

[3] S. Chambon, M. N. Galtier, P. J. Arnal, G. Wainrib, and A. Gramfort, “A deep learning architecture for temporal sleep stage classification using multivariate and multimodal time series,” arXiv, vol. 26, no. 4, pp. 758–769, 2017.

[4] B. Zhai, I. Perez-Pozuelo, E. A. D. Clifton, J. Palotti, and Y. Guan, “Making Sense of Sleep: Multimodal Sleep Stage Classification in a Large, Diverse Population Using Movement and Cardiac Sensing,” Proc. ACM Interactive, Mobile, Wearable Ubiquitous Technol., vol. 4, no. 2, 2020, DOI: 10.1145/3397325.

[5] O. Tsinalis, P. M. Matthews, and Y. Guo, “Automatic Sleep Stage Scoring Using Time-Frequency Analysis and Stacked Sparse Autoencoders,” Ann. Biomed. Eng., vol. 44, no. 5, pp. 1587–1597, 2016, DOI: 10.1007/s10439-015-1444-y.

[6] S. Mousavi, F. Afghah, and U. Rajendra Acharya, “SleepEEGNet: Automated Sleep Stage Scoring with Sequence to Sequence Deep Learning Approach,” arXiv, pp. 1–15, 2019, DOI: 10.13026/C2C30J.

[7] A. Vilamala, K. H. Madsen, and L. K. Hansen, “Deep convolutional neural networks for interpretable analysis of EEG sleep stage scoring,” IEEE Int. Work. Mach. Learn. Signal Process. MLSP, vol. 2017-Septe, no. 659860, pp. 1–6, 2017, DOI: 10.1109/MLSP.2017.8168133.

[8] O. Tsinalis, P. M. Matthews, Y. Guo, and S. Zafeiriou, “Automatic Sleep Stage Scoring with Single-Channel EEG Using Convolutional Neural Networks,” arXiv, 2016, [Online]. Available: http://arxiv.org/abs/1610.01683.

[9] S. Chambon, M. N. Galtier, P. J. Arnal, G. Wainrib, and A. Gramfort, “A deep learning architecture for temporal sleep stage classification using multivariate and multimodal time series,” IEEE Trans. Neural Syst. Rehabil. Eng., vol. 26, no. 4, pp. 758–769, 2018.

[10] S. Pathak, C. Lu, S. B. Nagaraj, M. van Putten, and C. Seifert, “STQS: Interpretable multi-modal Spatial-Temporal-seQuential model for automatic Sleep scoring,” Artif. Intell. Med., vol. 114, no. January, p. 102038, 2021, DOI: 10.1016/j.artmed.2021.102038.

[11] J. Lin, S. Pan, C. S. Lee, and S. Oviatt, “An Explainable Deep Fusion Network for Affect Recognition Using Physiological Signals,” in Proceedings of the 28th ACM International Conference on Information and Knowledge Management, 2019, pp. 2069–2072, DOI: https://doi.org/10.1145/3357384.3358160.

[12] C. Molnar, Interpretable Machine Learning A Guide for Making Black Box Models Explainable, 2018th-08–14th ed. Lean Pub, 2018.

[13] M. Youness, “CVxTz/EEG\_classification: v1.0,” 2020. https://github.com/CVxTz/EEG_classification (accessed Jan. 05, 2021).

[14] G. Al et al., “PhysioBank, PhysioToolkit, and PhysioNet: Components of a New Research Resource for Complex Physiologic Signals,” Circulation, vol. 101, no. 23, pp. e215–e220, 2000, [Online]. Available: http://circ.ahajournals.org/content/101/23/e215.full.

[15] B. Kemp, A. H. Zwinderman, B. Tuk, H. A. C. Kamphuisen, and J. J. L. Oberye, “Analysis of a sleep-dependent neuronal feedback loop: the slow-wave microcontinuity of the EEG,” IEEE Trans. Biomed. Eng., vol. 47, no. 9, pp. 1185–1194, 2000, DOI: 10.1109/10.867928.

[16] A. Swetapadma, “Novel approach for sleep disorder monitoring using a finite-state machine for localities lacking specialist physicians,” IET Sci. Meas. Technol., vol. 11, no. 8, pp. 1099–1103, 2017.

[17] C. Iber, S. Ancoli-Israel, A. L. Chesson, and S. F. Quan, “The AASM Manual for Scoring of Sleep and Associated Events: Rules, Terminology, and Technical Specifications.” 2007.

[18] D. P. Kingma and J. Ba, “Adam: A method for stochastic optimization,” arXiv Prepr. arXiv1412.6980, 2014.

